# Does a passerine hibernate?

**DOI:** 10.1101/808097

**Authors:** Brian K. McNab, Kerry Weston

## Abstract

The thermal physiology of the highly endangered Rock Wren (*Xenicus gilviventris*) from New Zealand is examined. It is a member of the Acanthisittidae, a family unique to New Zealand. This family, derived from Gondwana, is thought to be the sister taxon to all other passerines. Rock Wrens permanently reside above the climatic timberline at altitudes from 1,000 to 2,900 meters in the mountains of South Island. They feed on invertebrates and in winter face ambient temperatures well below freezing and deep deposits of snow. Their body temperature and rate of metabolism are highly variable. Rock Wrens regulate body temperature at ca. 36C, which in one individual decreased to 33.1C at an ambient temperature of 9.4C, which returned to 36C at 30.1C; its rate of metabolism decreased by 30%. The rate of metabolism in a second individual twice decreased by 35%, nearly to the basal rate expected from mass. The Rock Wren food habits, entrance into torpor, and continuous residence in a thermally demanding environment suggest that it may hibernate. For that conclusion to be accepted, evidence of its use of torpor for extended periods is required. Those data are not presently available. Acanthisittids are distinguished from other passerines by the combination of their temperate distribution, thermal flexibility, and a propensity to evolve a flightless condition. These characteristics may reflect their phylogenetic status, but they are so different from those found in other passerines that it is more likely that they reflect the geographical isolation of acanthisittids in a temperate environment for 85 million years in the absence of mammalian predators.

## INTRODUCTION

The question whether a passerine hibernates raises the broader question whether any bird hibernates. Commonly accepted is that the Poor-will (*Phalaenoptilus nuttallii*) hibernates based on observations (Jaeger, 1948, 1949) 70 years ago of torpid individuals found along a canyon wall in southern California in the same place several times. Poor-wills, like other caprimulgids, readily enter torpor (Brauner, 1952; Marshall, 1955; Bartholomew et al., 1957, 1963; Howell and Bartholomew, 1959; Ligon, 1970), but that does not guarantee that they hibernate.

Confusion between torpor and hibernation often occurs (McNab and O’Donnell, 2018). All examples of torpor do not represent hibernation, although hibernation is based on torpor. Hummingbirds in the tropics do not hibernate, even though they usually go into torpor at night reflecting a day of intense flight and a small mass. Hibernation is a behavior characterized by an extended period of continuous torpor in winter. (L. *hibernus*, winter.) Short-term torpor and hibernation represent extremes along a temporal continuum. The existence of an extended period of torpor is the basis for judging whether Poor-wills hibernate. Three Poor-wills were fed and kept from fall to spring in a large shed, after which they were released (Marshall, 1955). They were exposed to ambient temperatures from −5 to 22.5°C. The longest period of continuous torpor in an individual was four days. One was said to have “…hibernated every morning…” and “…a bird in deep hibernation at dawn would have been active through the previous evening before midnight “ (p. 132). This is not hibernation; it is short-term torpor. Poor-wills may hibernate, but there is no conclusive evidence in the form of an extended period of continuous torpor that they do.

A clear example of the temporal continuum of torpor is found in a small (7 g), solitary, insectivorous bat, the eastern pipistrelle (*Perimyotis subflavus*). It is distributed in eastern North America from southern Canada to central Florida. In Kentucky, half of the torpid individuals awoke after *ca.* 40 days of continuous torpor at a cave temperature of 10°C (Davis, 1965). In northern Florida half became active after 4 days of continuous torpor at a cave temperature of 16°C (McNab, 1974). This bat hibernates in the northern latitudes of its distribution, but uses short-term torpor in Florida because of the temperature dependence of torpor. The conversion of hibernation to short-term torpor in this species reflects latitude. In contrast, some temperate bats are committed to hibernation because copulation occurs in the fall and winter and therefore females must remain torpid to delay pregnancy until spring. An example is *Myotis grisecesens*, females of which in fall migrate from northern Florida to caves in Alabama that permit extended periods of torpor (McNab, 1974). This is required because cave temperatures in northern Florida were as high as 14°C; this species is larger (8-12 g) than the pipistrelle and hibernates in clusters, which requires even lower cave temperatures than for solitary species. Thus, some endotherms are committed to hibernation, whereas others have a flexible approach to ambient temperature.

The suggestion that a passerine may hibernate is completely unexpected. A few passerines go into torpor (McKechnie and Lovegrove, 2002; Schleucher, 2004), principally small, tropical insectivores and frugivores. Temperate swallows enter torpor (Lasiewski and Thompson, 1966; Serventy, 1970; Prinzinger and Siedle, 1988) in response to the unreliability of flying insects as food, but avoid harsh winters through migration to warm temperate and tropical environments. A similar behavior is found in temperate swifts and hummingbirds. In contrast, the Rifleman (*Acanthisitta chloris*), a sedentary passerine endemic to New Zealand, has a flexible approach to cold conditions (McNab and Weston, 2018). It readily enters torpor at ambient temperatures between 10 and 25°C.

The body temperature of some cold-temperate passerines, especially tits and finches, can be forced to 34°C. That requires an exposure to ambient temperatures from −15 to −30°C (Steen, 1958; Reinertsen, 1983). For example, it took an exposure to −15°C for 3 to 4 hours, combined with inanition, to get body temperatures between 34 and 35°C in the Willow Tit (*Parus montanus*) (Reinertsen and Hafton, 1983, 1984), a combination of conditions unlikely to be encountered in its nocturnal shelters. Body temperatures of Black-capped Chickadees (*P. atricapillus*) decreased to 34°C, when exposed to an ambient temperature of 0°C for four to six hours, but they were unable to arouse to their normal body temperatures at room temperature (Chaplin, 1976). These conditions are not required by the Rifleman to enter torpor. Furthermore, “[w]hen the [tits and finches] had become acclimated to constant cold (− 10°C) and were supplied with plenty of food, none entered into a hypothermic state” (Reinertsen, 1983, p. 276).

The highly endangered Rock Wren (*Xenicus gilviventris*) and the Rifleman are the only surviving members of the Acanthisittidae, the New Zealand ‘wrens.’ This passerine family is considered to be the sister taxon to all other passerines (Hackett et al., 2008; Selvatti et al., 2015; Mitchell et al., 2016), a line derived from Gondwana (Ericson et al., 2002; Worthy et al., 2010). Acanthisittids are not related to the Northern Hemisphere wrens (Troglodytidae), which are Oscine passerines. Of the eight species of acanthisittids, four of the six extinct species were flightless, the evolution of a flightless condition occurring at least three times (Worthy et al., 2010; Mitchell et al., 2012). And the Rock Wren is a weak flier.

The Rock Wren lives above the climatically based timberline in the mountains of South Island. Its altitudinal distribution is from 1,000 to 2,900 m, where in winter it encounters very low ambient temperatures and several meters of snow. Active nests of Rock Wrens have been found buried within snow banks. This wren does not descend to lower altitudes in winter in spite of its food habits, which consist of invertebrates. How can this combination of characters and conditions be tolerated and to what extent are they reflective of their basal position of all passerines? A limited number of measurements on the thermal biology of Rock Wrens were made, the results of which are reported here.

## MATERIAL AND METHODS

### Animals

Rock Wrens were captured at the Homer-Gertrude Cirque (*ca.* 1,000 m a.s.l.; 44.76 °S, 168.00 °E), Fiordland National Park, South Island, New Zealand. The habitat is comprised of extensive boulder fields, rocky bluffs, and snow tussock (*Chionochloa* sp.). Birds were transferred to the nearby Knobs Flat Research Station, in the lower Eglinton Valley, Fiordland National Park.

### Methods

Measurements of energy expenditure were made in the laboratory between 19:00 and 01:00 h, 5–7 h after capture when the wrens were inactive and post-absorptive. The oxygen consumption of the wrens was measured when contained in chambers that were placed in temperature-controlled containers. Two individuals were measured at the same time in separate chambers. Room air was drawn through the chambers, the exiting air scrubbed of carbon dioxide and water. The flow rate of the air stream was measured by a TSI 4140 flow meter (TSI Instruments Ltd, UK). It corrected the air volume to standard conditions of pressure (760 mm Hg) and temperature (0°C). The oxygen content of the air exiting the flow meter was measured by a S-3A/II Applied Electrochemistry oxygen analyzer (USA), its electrical outputs sent to a NGI strip-chart recorder (Austria). Measurements usually lasted for 1.5 to 2 hours or until a steady-state oxygen concentration was obtained. Three or four measurements, each at a different temperature, were made on each individual in a night. At the end of each temperature exposure, cloacal temperature and body mass were measured. The morning after measurement, the birds were released at their place of capture.

## RESULTS

Measurements of body temperature and rate of metabolism were made on six Rock Wrens. Mean body mass was 15.3 ± 0.23 g (n = 21). A very large variation occurred in body temperature (Fig. 1a). Whereas most cool-to cold-temperate passerines have body temperatures during their resting period between 39 and 42°C (McNab, 1966), in the Rock Wren it was usually between 36 and 37°C. Individual #1 had a body temperature of 36.5°C at ambient temperatures between 20 and 30°C (Fig. 1a). At an ambient temperature of 9.4°C, its temperature decreased to 33.1°C from which it spontaneously returned to 36.0°C within a half hour when exposed to 30.1°C. The high variability in body temperature in this species does not represent a failure of temperature regulation. A failure is illustrated by a decrease of body temperature with a decrease in ambient temperature, which is not seen here, an usually an inability to arise from torpor. Some of the variability is associated with activity, as in individual #2 (Fig. 1a). The variation in body temperature makes it difficult to estimate the Rock Wren’s regulated body temperature. At ambient temperatures between 12 and 30°C, an estimate of mean body temperature is 36.4 ± 0.15°C (n = 10).

**Fig. 1.**
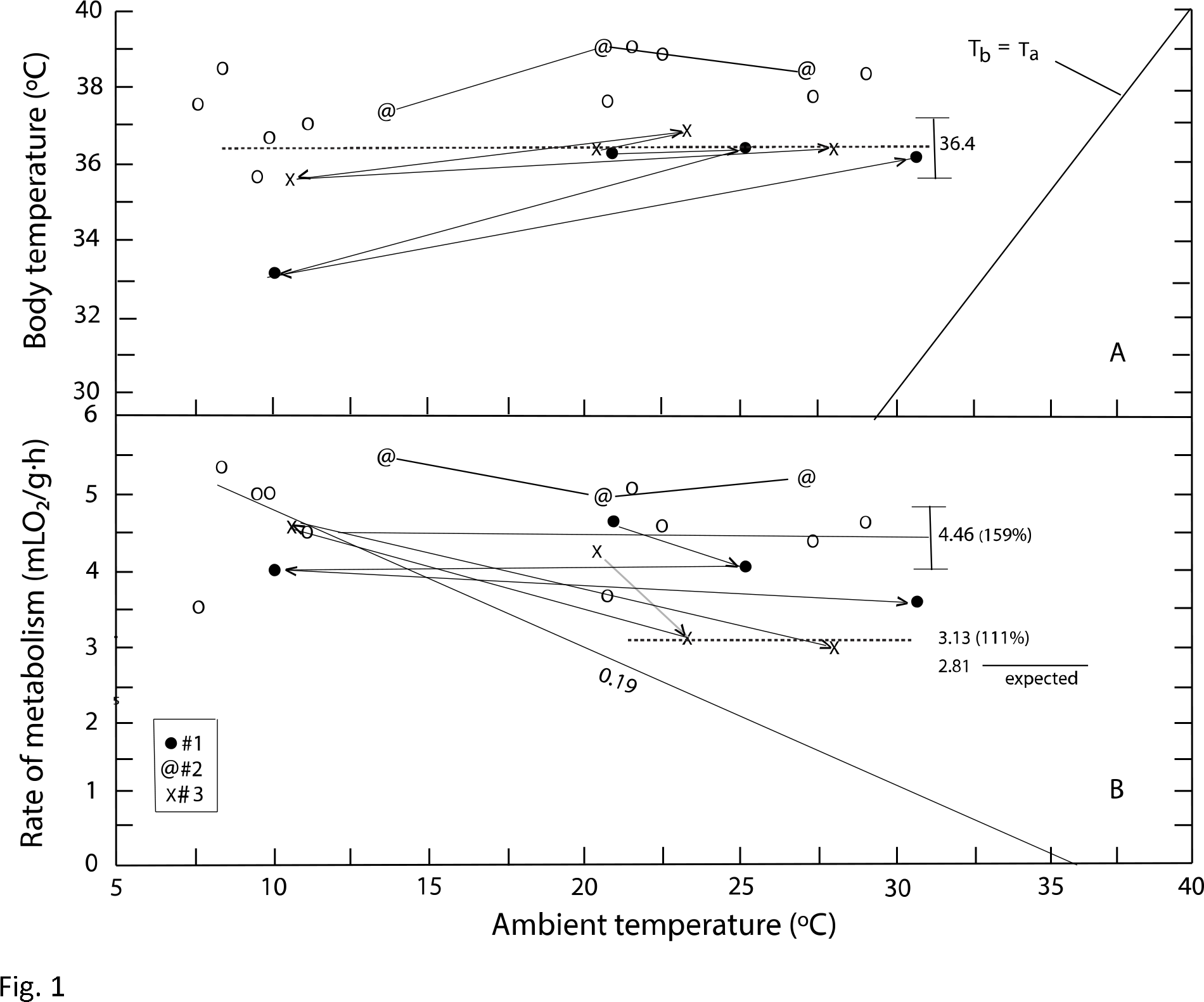
(**a**) The body temperature of six Rock Wrens (*Xenicus gilviventris*) as a function of ambient temperature. (**b**) Rate of metabolism as a function of ambient temperature.

As expected from the variation in body temperature, rate of metabolism is highly variable (Fig. 1b). Individual #2 had a high, variable rate of metabolism, as expected from its high, variable body temperature. The variability of rates in this species is so great that it is very difficult to estimate a basal rate of metabolism. A mean rate at ambient temperatures between 20 and 30°C is 4.46 ± 0.07 mLO_2_/g•h (n = 9); however, this mean is 159% of the basal rate expected from mean mass, 2.81 mLO_2_/g•h (McNab, 2009). This is unlikely to be a good estimate of basal rate, especially as several individuals had lower rates. One individual (#3) had two measurements that decreased by about 35% to 3.01 and 3.25 mLO_2_/g•h, the mean of which is 3.13 mLO_2_/g•h), which is 111% of the value expected from mass (Fig. 1b). These values were not accompanied with a reduction in body temperature. Whether this is a good estimate of basal rate is unclear, but it is more likely than the first estimate. Thermal conductance equals 0.19 mLO_2_/g•h•°C, which is 112% of the value expected from mass, about standard for its mass (Aschoff, 1981).

The detailed pattern of body temperature and rate of metabolism in individual #1 is described as a function of ambient temperature and time (Fig. 2). Its body temperature usually was ca. 36.5°C. But when exposed to 9.4°C, its immediate response was an increase in its rate of metabolism to compensate for the increased temperature differential with the chamber, after which the rate decreased by *ca*. 30% and body temperature to 33.1°C. The decrease in rate continued with a time lag when exposed to 30.1°C. At the end of the experiment, body temperature had increased to 36.0°C. If the exposure to 9.4°C had lasted longer, body temperature and rate of metabolism might have decreased further, but it had been exposed to the low ambient temperature for ca. 1.5 hours.

**Fig. 2.**
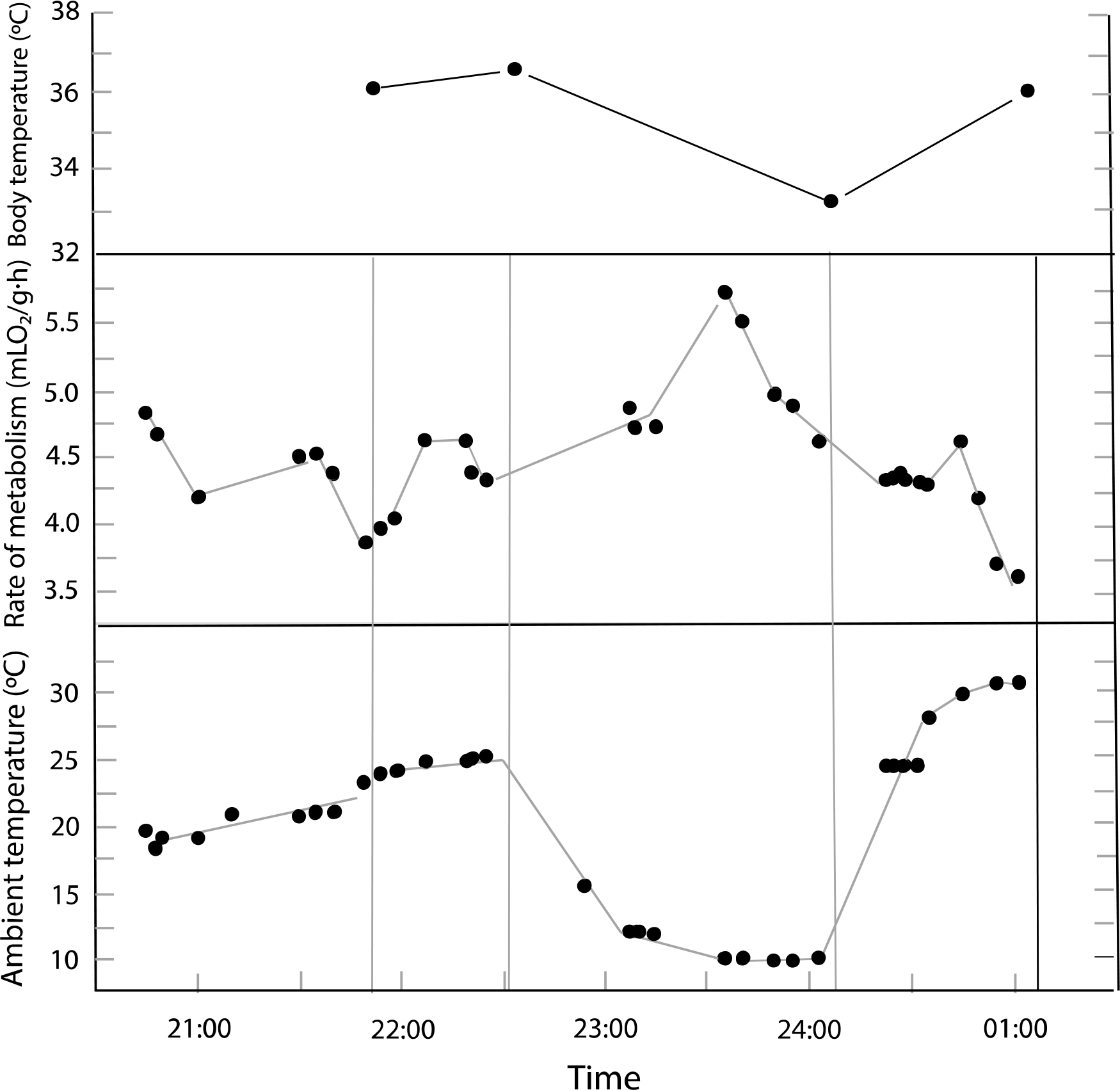
Body temperature, rate of metabolism, and ambient temperature as a function of time in individual #1.

## DISCUSSION

The Rock Wren has a highly variable body temperature and rate of metabolism at all ambient temperatures between 10 and 33°C. The variability is such that it is difficult to define the characteristics that are usually used to describe the energetics of an endotherm, a regulated body temperature and basal rate of metabolism. A similar a condition occurs in the Rifleman (McNab and Weston, 2018), its only living relative. What is striking is the propensity of the Rock Wren to have such a low, variable body temperature and rate of metabolism, while having a somewhat high basal rate, although its level is not clearly defined. The wren’s entrance into torpor is a regulated state, as it is in the Rifleman, which is demonstrated by their ability to spontaneously arise to their normal body temperatures. The two living members of this family have a pattern in energetics that is distinctive in passerines committed to residency in a temperate environment.

The behavior of these two species is very different from what occurred in the finches and tits (Reinertsen, 1983). Unlike the Rock Wren, the Rifleman is found at altitudes from sea level to 1,000 m in forested environments. It gleans an insect diet from surfaces, especially tree trunks, cool weather not likely disrupting its food supply for an appreciable period. There is no evidence to suspect that the Rifleman hibernates, although measurements of nest temperatures in winter may clarify that possibility.

The Rock Wren is committed to cold environmental conditions that it confronts at altitudes > 1,000 m in the mountains of South Island in winter. The environmental conditions that the Rock Wren faces, in combination with its thermally vulnerable food habits, entrance into torpor, and sedentary lifestyle, make it a likely candidate for hibernation. These characteristics differ from those of finches and tits, which are granivorous, a food supply available throughout the year, unlike the invertebrate diet consumed by the Rock Wren.

Evidence of an extended period of torpor is required to conclude that the Rock Wren hibernates, which we think is likely. Given its highly endangered status, however, acquiring enough data will be difficult to obtain. The continuous direct or indirect measurements of body temperature in an occupied nest during winter may distinguish between the occurrence of short-term torpor and hibernation in the Rock Wren, as is also required to demonstrate hibernation in the Poor-will.

What is clear is that the acanthisittids are physiologically distinctive with thermal behavior unknown in any other temperate passerine family, especially when coupled with their propensity to evolve a flightless condition. The extinct members of the family probably were also thermally flexible, given the behavior of their two living relatives, their small masses, and flightless status. The extent to which these characteristics emerged from the family’s unique phylogenetic position is unclear, unless it reflects a thermal flexibility not found in other temperate passerines. However, this flexibility is more likely to be a response to residence on a temperate landmass that has been independent of Gondwana for over 85 million years in the absence of mammalian predators. These conditions are unlikely to be present elsewhere, although the evolution of a flightless bunting (*Emberiza alcoveri*) occurred on the Canary Islands (Rando and López, 1999), reflective of a widespread tendency on islands (Wright et al. 2016), which have had the same protection from predators as New Zealand had.

A question has existed whether New Zealand was completely submerged some 25 mya, 60 million years after its separation from Gondwana. The occurrence of this family and many other iconic groups in the Early Miocene suggests that some part of New Zealand was not submerged and that this family reflects a vestige of a Gondwanan ancestry (Worthy et al., 2010; Mitchell et al., 2012).

## Acknowledgements

We thank the staff of Fiordland Electrical for assistance in setting up electronic equipment, and staff at the Knobs Flat Research Station, particularly Jo Carpenter for assistance in capturing the birds. We appreciate a review of the manuscript and suggestions by David Steadman.

